# Biomechanical effects of the addition of a precision constraint on a collective load carriage task

**DOI:** 10.1101/2021.07.09.451742

**Authors:** Nour Sghaier, Guillaume Fumery, Vincent Fourcassié, Nicolas A. Turpin, Pierre Moretto

## Abstract

Team lifting is a complex and collective motor task that possesses both motor and cognitive components. The purpose of this study was to investigate to what extent the biomechanics of a collective load carriage is affected when a dyad of individuals is performing a carrying task with an additional accuracy constraint. Ten dyads performed a first condition in which they collectively transported a load (CC), and a second one in which they transported the same load while maintaining a ball in a target position on its top (PC).

The recovery-rate, amplitude, and period of the center-of-mass (COM) trajectory were computed for the whole system (dyad + table = PACS). We analyzed the forces and moments exerted at each joint of the upper limbs of the subjects. We observed a decrease in the overall performance of the dyads when the Precision task was added, i.e., i) the velocity and amplitude of CoM_PACS_ decreased by 1,7% and 5,8%, respectively, ii) inter-subject variability of the Moment-Cost-Function decreased by 95% and recovery rate decreased by 19,2% during PC. A kinetic synergy analysis showed that the subjects reorganized their coordination in the PC.

Our results demonstrate that adding a precision task affects the economy of collective load carriage. Notwithstanding, the joint moments at the upper-limbs are better balanced and co-vary more across the paired subjects during the precision task. Our study results may find applications in domains such as Ergonomics, Robotics-developments, and Rehabilitation.

## 2 Introduction

Individual manual materials handling is commonly performed in many human activities. Its definition regroups essentially different tasks of load handling (i.e. the transport, support, lift, push, pull…) and has unfavorable ergonomic conditions (Directive 90/269/CEE). Different lifting techniques (Faber et al., 2007) and mechanical aids (Godwin et al., 2009) have been proposed to avoid the associated leisure. However, the most common alternative used is team lifting. It is, in appearance, a simple solution to carry a heavy load (i.e. too heavy to be carried safely by a single individual), a bulky object without mechanical aids. This strategy is then supposed to reduce the load on individuals performing the task (Sharp et al., 1997). It is also used in some sports such as Crossfit discipline when lifting “worm”, a heavy, long, and soft cylindrical bag (Claudino et al., 2018).

Furthermore, team lifting necessitates both motor and cognitive skills in order to control the movement coordination. For this reason, many authors consider it as a dual-task, e.g. workdays in ironwork (Faber et al., 2012). The tasks are called dual as they often undergo some interference, linked to a limited ability to share attention between the two task goals. Dual-task interference are commonly studied in psychology to highlight the cognitive limits of the human brain (Pashler, 1994). One well-known example of these cognitive limits is based on locomotion which can be used in interference paradigm (Yogev-Seligmann et al., 2008; Yogev-Seligmann et al., 2010). For example, walking along an L-shaped path while performing an arithmetic task deteriorates the mobility function. Any additional cognitive-task is considered as a limiting factor for the motor task since it induces a modification of gait pattern e.g., reduces gait speed, induces movement fluctuation and oscillation. Beach et al., (2006), showed that during a repetitive lifting task adding a precision placement challenges leads to an increase of lumbar spine load and an increase of the upper limb movement time.

Recent studies focused on lifting-Precision dual-task and showed that locomotor pattern was not affected when the subjects performed a dual-ask such as carrying a load (20% of mean body mass) on the shoulders (Castillo et al., 2014), or carrying a load (21% and 36 % of mean body mass) on the back(Ackerman and Seipel, 2014; Bastien et al., 2016). However, other studies reported a walking pattern affected by the dual-task when the load is balanced on the top of the head (Heglund et al., 1995) or when it is carried collectively (Fumery et al., 2018). In certain jobs, a collective load transfer is simultaneously associated to a precision task (e.g. talking with the patient in nurses, administrating medication in stretchers bearer, Industrial manufacturing in workman). This occurs especially in paramedics or during search and rescue activities (Gamble et al., 1991; Restorff, 2000; Barnekow-Bergkvist et al., 2004; Leyk et al., 2007).

Despite this practical relevance, only few studies deal with biomechanical aspects of collective transport of a load. However, it is essential to conceive aids, exoskeletons and collaborative robots dedicated to assisting humans in such a task. Fumery et al., (2018) studied the energetic exchanges during a collective load carriage to investigate whether two individuals transporting an object behave economically. The authors showed that the external energetic exchanges occurring during this type of transport was as efficient as those occurring in single gait when the load is below 10% of the total body mass of the dyad. In our study, we reproduce the protocol of Fumery et al. to investigate the locomotor pattern of ten paired individuals carrying a box collectively and to compare these walking patterns when this task is performed simultaneously with a precision task. This precision task consists in maintaining a ball in the center of a circular target drawn on the top of the box. Thanks to handle sensors and kinematic data, we also record the forces and moments applied on the box then compute the constraints at the arms and back joints using the inverse dynamics bottom-up procedure. The purpose of this study is to investigate to what extent the performance of a collective load transport is impacted when a cognitive task is performed simultaneously, and how the control of the cognitive task is shared across the subjects. As it has been observed in single subjects walking and performing a cognitive task, we hypothesize that gait performance of the individuals walking while transporting collectively a load is disturbed when they perform a precision task requiring precision. Fumery et al. (2018) demonstrating that the pattern of a dyad carrying a light load was not affected by the collective task, our hypothesis is that, using the same load ratio, the precision task is the single factor affecting the walking-carrying pattern of the paired subjects.

## 3 Materials and Methods

### 3.1 Population

Ten pairs of healthy male individuals (mean±s.d.: volunteer 1 - at the left side of the load: height = 1.77±0.07 m, mass = 74.78±9.00 kg; volunteer 2 - at the right side of the load: height = 1.77±0.05 m, mass = 74.54±12.38 kg) participated in the experiments. The individuals had no orthopedic disabilities, no dysfunctions of the locomotor system, no neurological or vestibular diseases, no visual deficits and no proprioceptive disorders or dementia.

This study was carried out in accordance with the requirement of a non-interventional study given by the CNRS bioethical office. The study was approved by the Research Ethics Committee of the University of Toulouse, France (number IRB00011835-2019-11-26-172, Université Fédérale de Toulouse IRB #1). All subjects gave verbal and written informed consent in accordance with the Declaration of Helsinki.

### 3.2 Experimental Protocol

Two conditions were performed by the subjects. In the first condition (Control Condition: CC) they walked side by side at spontaneous speed while carrying a box (mass = 13.41 kg, size: 0.40×0.40×0.28 m) equipped with two lateral handle sensors (Sensix, France). The mass of the box plus the sensors was 14.250 kg thus almost 10% of the body-mass of the two volunteers. In order to get accustomed to the task the subjects performed three successive trials. Only the last and third trial was retained for the analysis.

In the second condition (Precision Condition: PC), the individuals were instructed to transport the box, while performing an accuracy task consisting in keeping a ball (diameter: 19mm, mass: 2g) in the center of a circular target drawn on the top of the box (Fig 1). Subjects were not allowed to orally communicate during the experiments.

**Figure 1.**
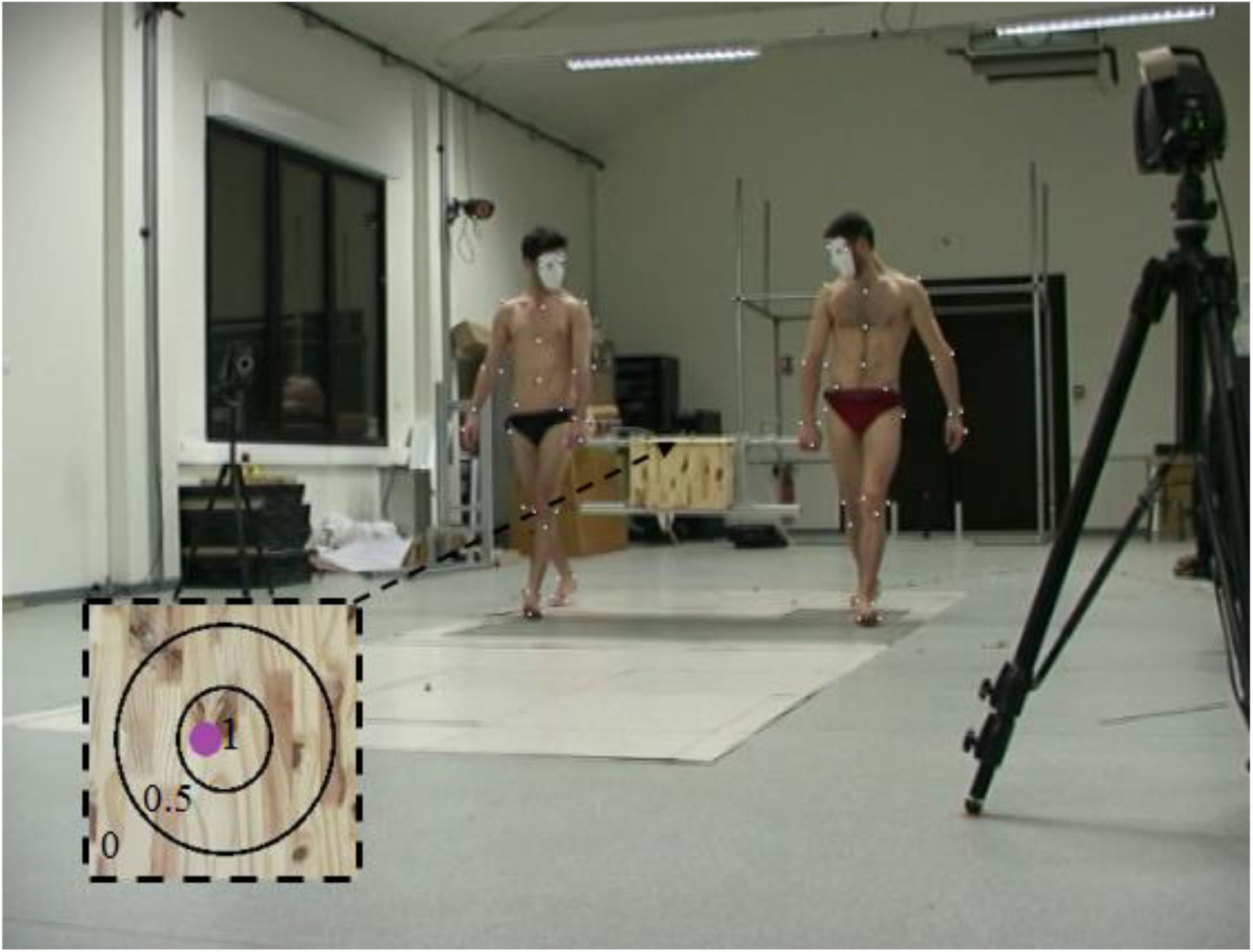
Experimental setup: Collective load carriage performed with a precision task (PC). The dyad carried a box (mass = 13.41 kg, size: 0.40x 0.40 × 0.28) while maintaining a ball (diameter=19mm, mass=2g) in the center of a targeted position (diameter of the small and large circle: 120 mm and 240 mm respectively) on its top. If the ball was maintained in the small circle, the dyad obtained a 1-point Scorep. If it was maintained between the small and large circle, then they obtained 0,5-point Scorep. Else, they obtained 0-point Scorep.

### 3.3 Kinematic and Kinetic Data Acquisition

Motion capture data were collected using thirteen infrareds (11 MX3 and 2 TS40) transmitter-receiver video cameras (Vicon^©^, Oxford metric’s, Oxford, United Kingdom) sampled at 200 Hz. Forty-two retro-reflective markers were placed on bony landmarks and on the navel of each subject (according to Wu et al., 2002, 2005) and fourteen on the box. The ball used during the PC tests was reflective as well and was tracked by the Vicon^©^ system.

In order to record the gait pattern at constant speed (i.e. to exclude the acceleration and deceleration phases at the beginning and end of each trial) the volume calibrated by the Vicon^©^ system (30 m^3^) was located in the middle of the 20m-long walkway crossed by the subjects. The reflective marks were tracked to define the kinematics of the Poly-Articulated Collective System (PACS) formed by the two individuals and the load they carry (Zatsiorsky, 1983; Moretto et al., 2016). The data were recorded on one gait cycle defined by the first heel strike of the first subject and the third heel strike of the second subject of the PACS to ensure a cycle of each subject. The 3D reconstruction was performed using Vicon Nexus 1.8.5^©^ software.

The two lateral handles used to transport the box were equipped with Sensix^®^ force sensors sampled at 2000 Hz. A 4^th^ order Butterworth filter and a 5 Hz and 10 Hz cut frequency have been applied to analyze the positions of the markers and the forces exerted on the box handles, respectively.

### 3.4 Computed parameters

#### 3.4.1 Trajectory of the CoM_PACS_

The De Leva Anthropometric tables (de Leva, 1996) was used to estimate the mass m_i_ and the CoM of each segment i (CoM_i_) of the PACS and to compute its global CoM (CoM_PACS_) as follow:

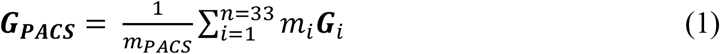

with ***G***_***PACS***_ the 3D position of the CoM_PACS_ in the frame R (the global coordinate system), *m*_*PACS*_ the mass of the PACS, *n* the number of PACS segments (i.e. 16 segments per volunteer plus one segment for the box) and ***G***_***i***_ the 3D position of the CoM_i_ in the frame R. The CoM of the box was determined at the intersection point of the vertical lines obtained by hanging it with a thread fixed at different positions. The material used for the box construction, i.e. wood and aluminium, was considered as not deformable.

According to Holt et al., (2003), the amplitude (*A* = *Z*_max_ – *Z*_min_, with *Z* the height of the CoM_PACS_, in meters,) and the period (peak to peak, in percent of the gait cycle) of the CoM_PACS_ were also assessed.

The forward kinetic (*W*_kf_), as well as the vertical (*W*_v_) and external work (*W*_ext_) of the CoM_PACS_ were computed according to the method of Bastien et al. (2016). Then based on the external work, the percentage of energy recovered of the CoM_PACS_ in the sagittal plane was computed (called recovery rate *RR* in Fumery et al., 2018a, 2018b). This parameter assess the amount of energy transferred between the potential and the kinetic energy (Eqn 2).

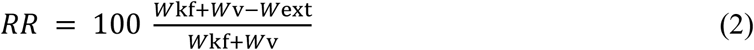

The closer the value of *RR* to 100%, the more consistent the locomotor pattern is with the inverted pendulum system (IPS) model of locomotion (Cavagna et al., 1963; Willems et al., 1995; Gomeñuka et al., 2014; Bastien et al., 2016). In this study, the trajectory of CoM_PACS_ and CoM of an inverted pendulum have been investigated.

#### 3.4.2 Forces and moments at the joints of the upper limbs

Sensix force sensors recorded the forces and moments applied by each individual on the two box handles. Before the computation, the data of the sensors located by specific markers were transfer to the Galilean frame of the laboratory using rotation matrix. A cross correlation method has been applied in order to analyze the coordination between the forces produced by both subjects. To investigate whether the movement of the box results from an action-reaction strategy, we computed the time lag required for the position of the left side and right side of the box to be the same on the medio-lateral, antero-posterior and vertical axis in CC and PC. The coordination was assessed through the forces exerted on three directions (medio-lateral, antero-posterior and vertical axis). This results will reflect the level of coordination of two subjects during a collective transport

In order to quantify muscular constraints produced at the upper limb, the Inverse Dynamic Method was used to estimate forces and moments at each joint of the upper limb. The Moment Cost Function was then computed (kg.m^2^.s^-2^, Costes et al., 2018) as follow :

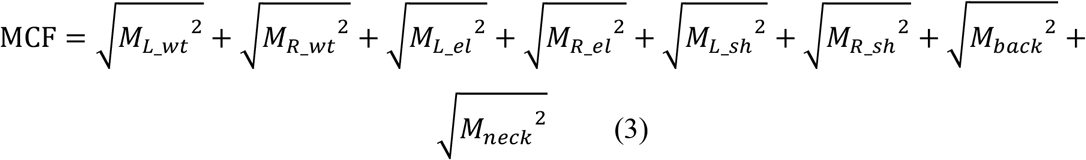

Where M_L_wt_, M_R_wt_, M_L_el_, M_R_el_, M_L_sh_, M_R_sh_, M_back_ and M_neck_ are the mean values over a PACS gait cycle of the three-dimensional left and right wrist, left and right elbow, left and right shoulder, top of the back and neck moments, respectively. 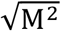 represents the Euclidian norm of M (i.e. 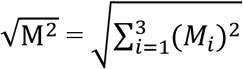, with M_i_ the i^-th^ component of the vector M).

Then, the MCF values of each individual was summed to obtain the total moment cost function (Total *MCF*). This Total *MCF* allows to quantify the global effort produced at the upper-limbs of the PACS during one gait cycle. Finally, the *MCF* difference (Δ *MCF*) was computed as the difference between the two individuals to investigate whether the subjects produced the same effort in the upper limbs during the load transport.

#### 3.4.3 Kinetic synergy analysis

We extracted the synergies by using a principal component analysis (PCA) applied to the wrist, elbow, shoulder, back, and neck joint moment on the right and left sides of the body. The PCA was used to reduce data dimensionality. It consisted in the eigen-decomposition of the co-variance matrix of the joint moment data (Matlab *eig* function). The joint moments data from one trial per condition were arranged in time × joint moment matrices. In this analysis we only used the y-component which is very close to the norm of the 3D joint moments, except that the y-component (medio-lateral) could be positive and negative. The joint moments were normalized by their amplitude and centered (mean removed) before application of the PCA. We called the eigenvectors extracted from the PCA, dynamic synergy vectors. We computed the VAF (Variance Accounted For) which corresponded to the cumulative sum of the eigenvalues, ordered from the greatest to the lowest value, normalized by the total variance computed as the sum of all eigenvalues. The synergy vectors retained were then rotated using a Varimax rotation method to improve interpretability.

We first extracted the synergy vectors for each experimental condition and each participant separately. In this analysis the initial data matrices were constituted of all available time frames in line, concatenated from one trial per condition, and of eight columns corresponding to each joint moment, namely the right wrist, left wrist, right elbow, left elbow, right shoulder, left shoulder, back, and neck. Based on a previous study we extracted 3 synergies in this analysis. We then performed a second analysis to identify possible co-variations between the joint moments of the two participants in each pair. The columns of the initial matrices were thus constituted of the joint moments of the two loaded arms, i.e., the right wrist, elbow, and shoulder joint moments of participant #1, plus the left wrist, elbow and shoulder joint moments of participant #2. Based on a previous study we extracted 2 synergies in this analysis. We used Pearson’s r to order the different synergies similarly between the different subjects and conditions.

#### 3.4.4 Accuracy score

A performance score (score_p_) was assigned to each image of the videos captured by the Vicon^©^ system (200 images/s). The score depended on the location of the ball in the target: 1 when the ball was inside the small circle, 0.5 when it was in-between the small and large circle and 0 when it was outside the large circle. The accuracy over the whole gait cycle was measured by an overall score (Score_accuracy_), expressed in percentage, and calculated as follows:

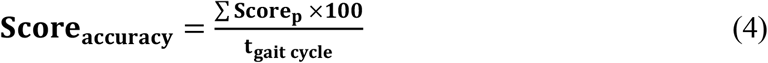

where t_gait cycle_ represents the number of Vicon^©^ images recorded along one gait cycle.

#### 3.4.5 Orientation of the upper part of the body

The head, shoulders and pelvis rotation angles were computed around the vertical axis of each individual in the two conditions. The angle was positive when the subjects turned towards the box they carried, otherwise it was negative. The distance between the forehead and the sternum (distance FOR-STE) was also computed in order to investigate the flexion of the cervical spine.

### 3.5 Data analysis

The data were analyzed with Matlab R2016b© and StatView 5.0© software. A paired t-test was used to compare the RRs, the amplitudes, the periods, the velocities of the vertical displacement of the CoMPACS, the head, shoulders and pelvis rotation angle and the length FOR-STE between the CC and PC condition. The significance threshold was set to 0.05. We computed with a cross-correlation method the time lag required for the position of the left side and right side of the box to be the same on the medio-lateral, antero-posterior and vertical axis in CC and PC.

We used the average subspace angles to compare the subspaces spanned by the synergy vectors (Knyazev and Argentati, 2002). In order to decide whether the subspaces were more similar than expected by chance, the confidence interval (CI) of random comparisons was computed. For this analysis, we generated pairs of random subspaces constituted each of either 3 unit vectors of dimension 8 (individual PCA analysis) or 2 unit vectors of dimension 6 (conjoint PCA analysis) and computed the mean subspace angle between them. The unit vectors were built using normally distributed pseudo-random numbers (Matlab randn function). We performed 10000 simulations in order to determine the 95%-CI of the mean subspace angle between the pairs of random subspaces. The confidence interval was 39.5°–70.0° (55.0±7.6°) for the individual PCA analysis and 36.3°–79.1° (57.7±10.7°) for the conjoint analysis.

We used Student tests for single mean to compare the subspaces angles to the lower bound CI with the assumption that similarity was higher than expected by chance when the angles were lower than the lower bound CI.

VAFs were compared with an ANOVA with one repeated measure (control vs. precision conditions) and one factor (participant #1 vs. participant #2) when synergies were extracted separately for each subject. For the conjoint analysis, a paired Student t-test was used. Subspace angles were compared with t-tests for dependent samples (paired t-test) when comparing the control and precision conditions and t-test for independent samples when comparing the two participants. Adjustments for multiple comparisons were performed by Bonferroni’s method. Initial level of significance was set to p<0.05.

## 4 Results

### 4.1 Dynamics analysis of the dual-task

#### 4.1.1 CoM trajectory

The CoM_PACS_ velocity significantly decreased from 1.40±0.14 m.s^-1^ in CC to 1.23±0.17 m.s^-1^ in PC (*t* =3.385, p=0.008). The CoM_PACS_ amplitude (Fig. 2A, *t*=3.704, *p*=0.005), significantly decreased from 2.87 ± 0.742 cm in CC to 2.29 ± 0.739 cm in PC (Table S3). However, the period of the CoM_PACS_ oscillation was not significantly affected by the precision task (Fig. 2B, *t*=0.842, p=0.422).

**Figure 2.**
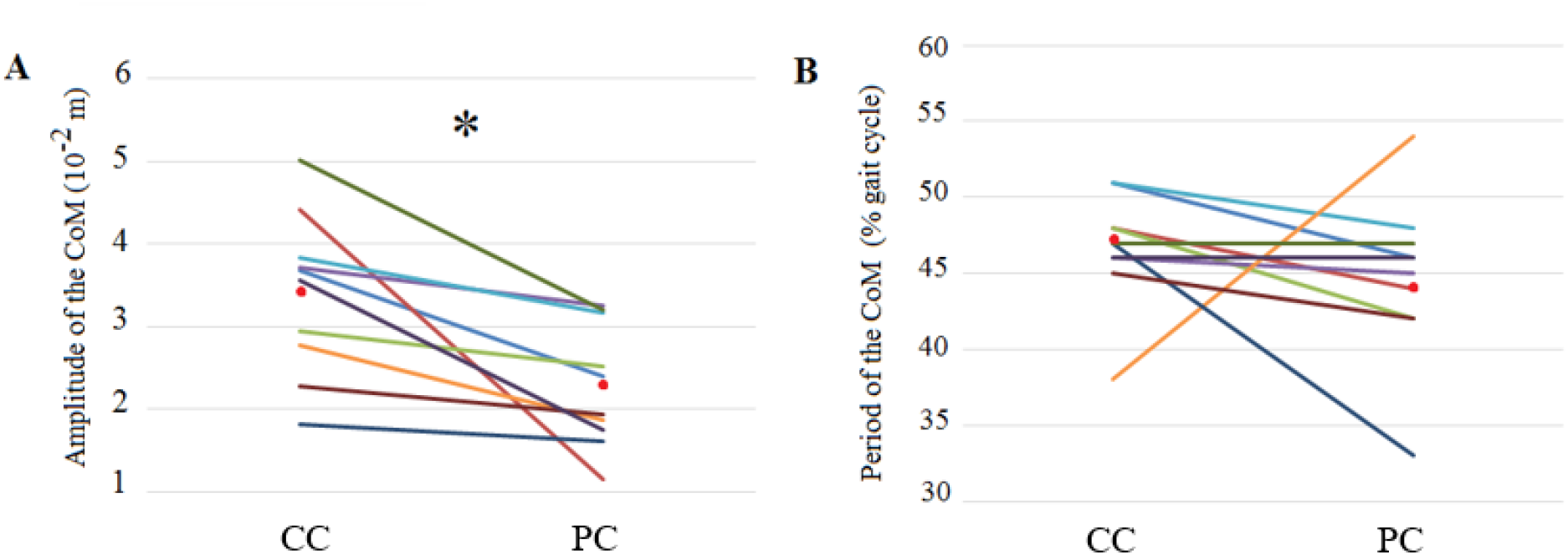
Amplitude (A) and period (B) of the vertical displacement of the CoMPACS in the Control Condition (CC) and the Precision Condition (PC). The mean value of each dyad (N=10) was computed for the CC and PC and linked. The red points represent the mean for each condition. The same color is assigned to each dyad in all figures. * = significant difference (p<0.05 paired t-test).

The percentage of energy recovered at the CoM_PACS_ significantly decreased between CC and PC (t=5.18, p<0.001) (Fig 3). This showed an alteration of the efficiency of the locomotor pattern of the dyad when the energy transfer between the potential and the kinetic energy was by 19,2% lower in PC compared to CC.

**Figure 3.**
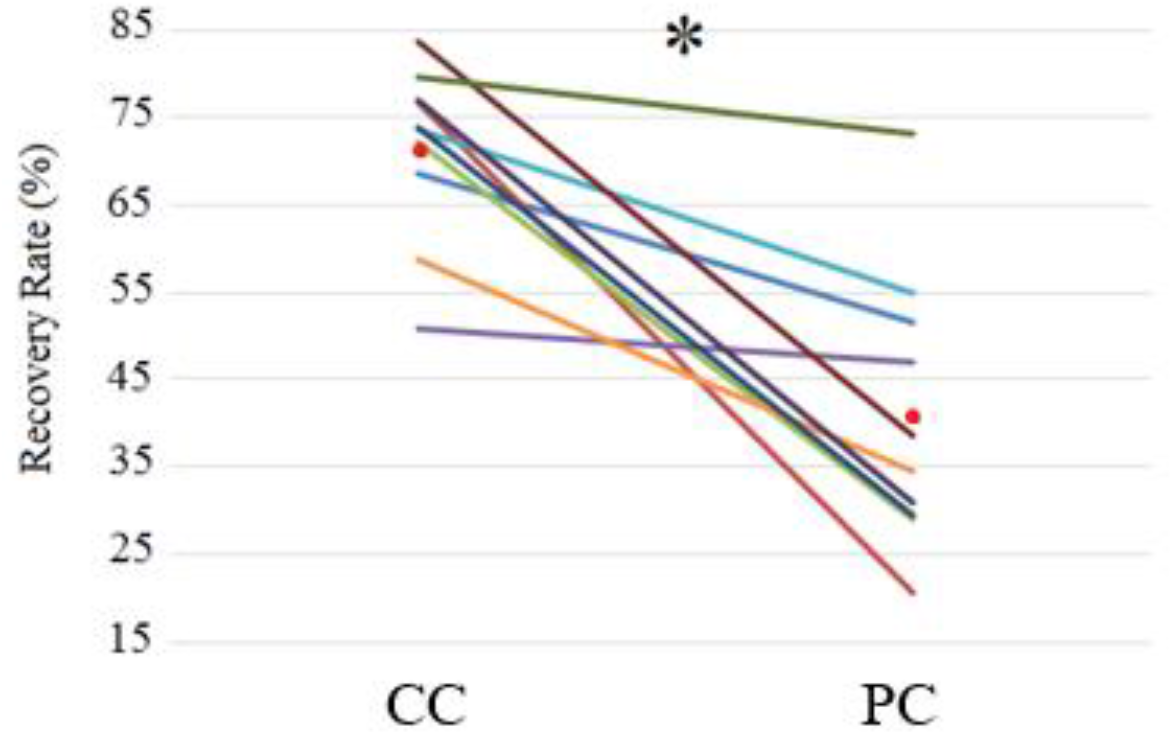
Recovery Rate (%) of each dyad (N=10) during Control Condition (CC) and Precision Condition (PC). The red points represent the mean for each condition. The same color is assigned to each dyad in all figures. * = significant difference (p<0.05 paired t-test).

#### 4.1.2 Forces applied to the handles

Significant differences were found between the correlation coefficients of the three components (x, y, z) of the forces applied to the handles by the subjects of the dyad for each condition. In the CC condition, R_Fx_ was lower than R_Fy_ (t = -3.45, p < 0,001) and R_Fz_ (t = -4.53, p < 0,01) and R_Fy_ was lower than R_Fz_ (t = -2.48, p = 0,04). In the PC condition, R_Fx_ was lower than R_Fy_ (t = - 6.06, p < 0,01) and R_Fz_ (t = -4.50, p < 0,01). However, no significant differences (Fig. 4A) were found between the CC and the PC conditions (t = -0.43, p = 0.675; t = -1.43, p = 0.188; t = - 1.02, p = 0.335 for the medio-lateral, antero-posterior and vertical axis, respectively) (Table S8).

**Figure 4.**
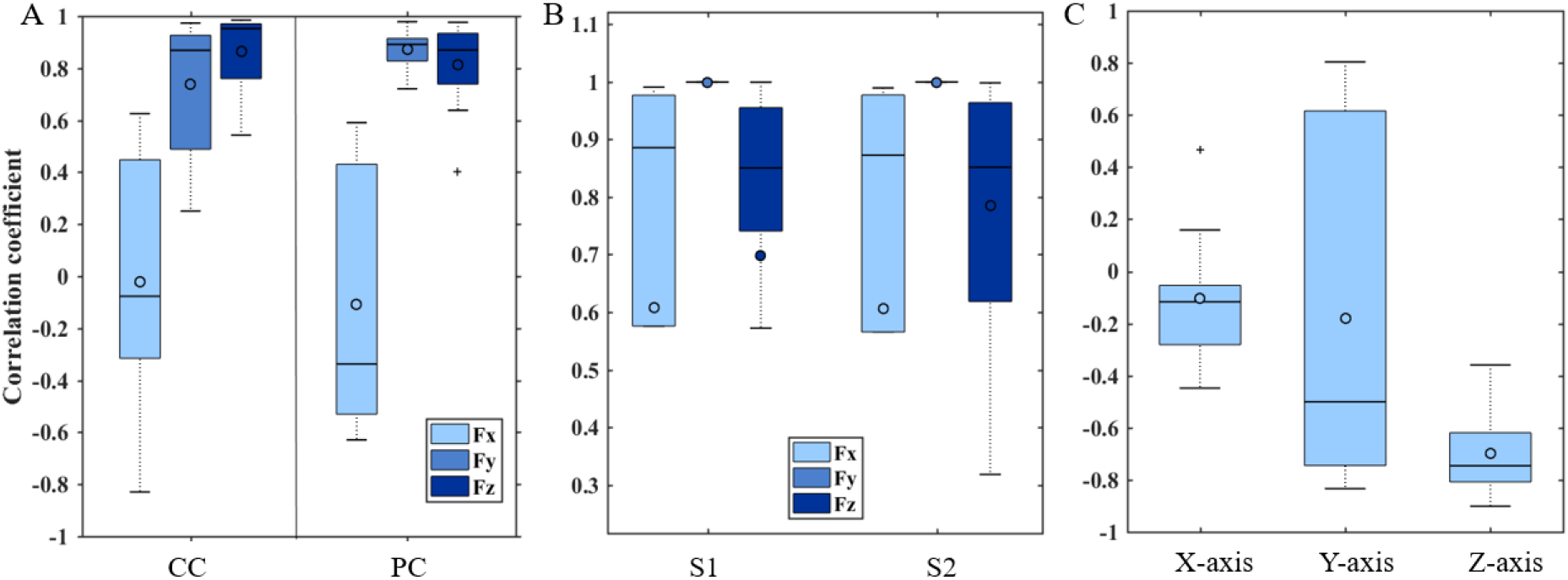
Boxes plot showing the distribution of the correlation coefficient (Coef). A : Coef of the forces produced by the individuals in each dyad on the box handles, on the medio-lateral (Fx), antero-posterior (Fy) and vertical axis (Fz) in the CC and PC conditions. B : Coef of the ball displacement and the handles displacement, on Fx, Fy, and Fz, in the CC and PC conditions. C : Coef of the ball trajectory and the sum of forces exerted by the subjects on the handles, on Fx, Fy, and Fz, in PC. N = 10 for each condition. * 0.05> p> 0.01; ** p <0.01 (paired Student t test). The upper horizontal line of the box represents the third quartile (75th percentile), the lower line of the box represent the first quartile (25th percentile), the middle value of the dataset is the median value (50th percentile) and the upper and lower horizontal lines outside the box represent respectively 90th percentile and 10th percentile. Cross-and circle represent respectively outlier and mean.

In the CC, the time lags were lower than 150 ms in the medio-lateral and antero-posterior axis and only one lag was higher than 150 ms for one dyad for the vertical axis (Table 1, lagZ 20 ± 50 ms). Concerning the PC, the time lags were also lower than 150 ms in the medio-lateral and antero-posterior axis and five dyads had a time lag higher than 150 ms for the vertical axis (Table 1, lagZ 180 ± 230 ms) (Table S7).

**Table 1.**
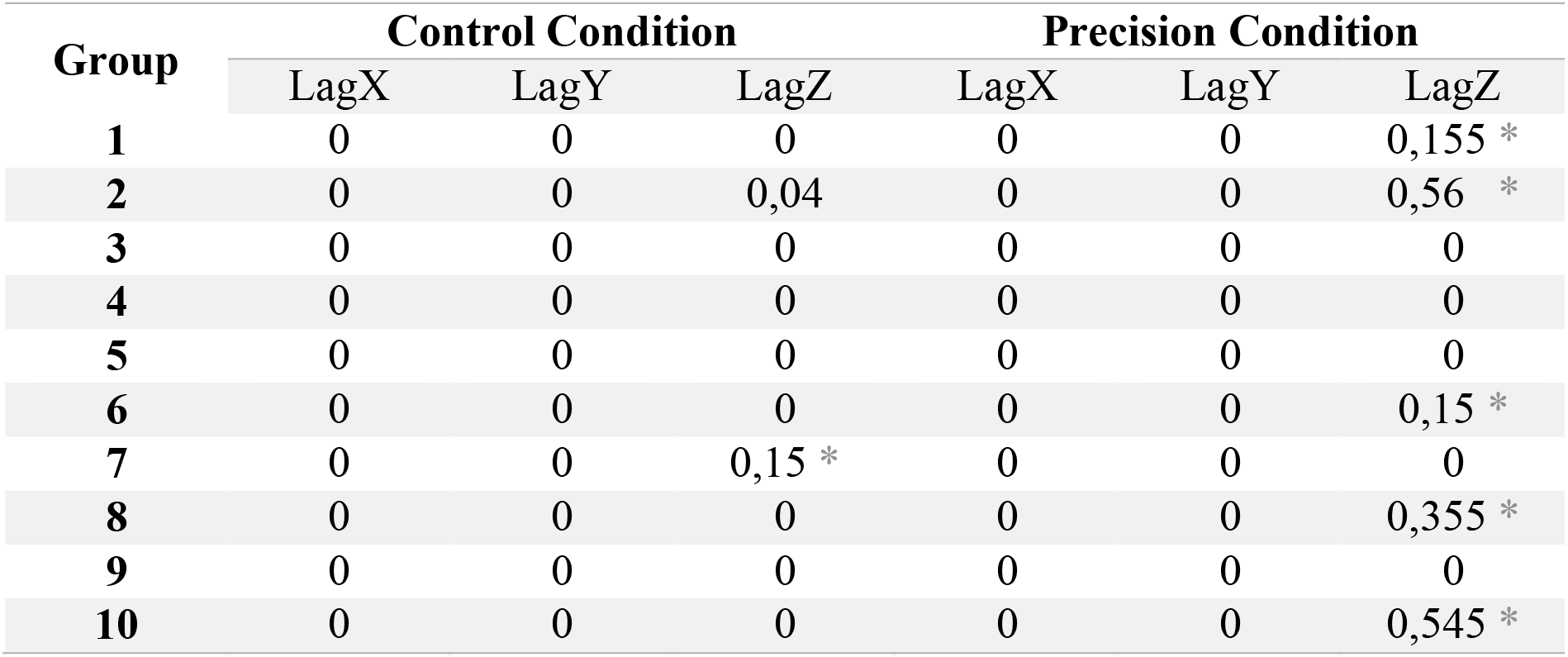
**The action-reaction strategy**, the time lag (s) required for the position of the left side and right side of the box to be the same on the medio-lateral, antero-posterior and vertical axis in CC and PC. **\* 150 ms < p (VanRullen and Thorpe,2001)**

#### 4.1.3 Kinetic synergy analysis

Consistent with a previous study we extracted 3 dynamic synergies for all subjects, which accounted for 96.3 ± 2.0 % of total variance on average (range [90.0–99.0]%). We found no effect of the side (being on the left or right side of the load) nor of the precision constraint on the VAF values (i.e., |ΔVAF|=1.9±0.4%, *p*-value = 0.24, η^2^=0.08 and |ΔVAF|=0.9±0.5%, *p*-value = 0.62, η^2^=0.01, respectively).

The dynamic synergies for each participant are depicted in Fig 5. The comparisons between participant #1 and participant #2 gave subspaces angles not different than expected by chance in the precision condition (i.e., 45.0±10.0° compared to 39.5°, Student t_9_=-1.34; *p*-value=0.21; Fig. 6A). For the other comparisons in Fig 6, subspace angles were lower than expected by chance (Fig 6, |Student t_9_ ≥7.6; *p*-value<0.001). The subspace angles were lower in the control condition than in the precision condition when comparing participant #1 and participant #2 (Fig. 6A, Cohen’s d= 1.9; Student t_9_=-4.95; *p*-value<0.001); showing an effect of the conditions on inter-subject similarities. The comparison between the control and precision conditions gave subspace angles of 27.2±5.7° on average with no differences between subjects (Fig. 6B, Cohen’s d= 0.61; Student t_9_=-1.93; *p*-value=0.09). The inter-condition subspace angles were lower than inter-subject subspace angles (i.e., 27.2±5.7° *vs*. 37.1±7.7°, Cohen’s d= 1.5; Student t_18_=3.30; *p*-value=0.004).

**Figure 5.**
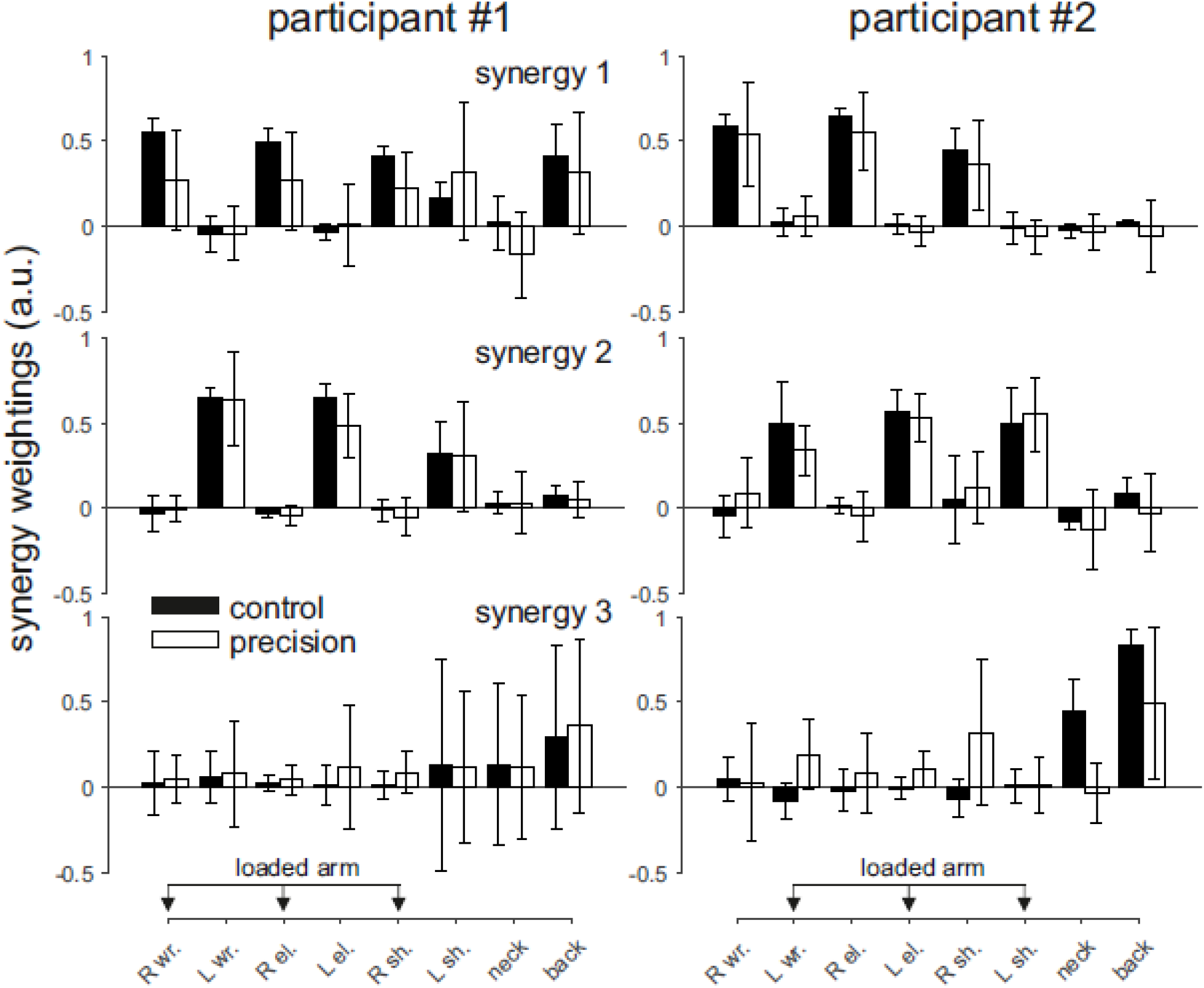
Dynamic synergy vectors. Three synergies accounted for more than 90.0% of total variance in all subjects and conditions. Participants labeled #1 were on the right side of the load and participants #2 on the left side. Wrist, elbow and shoulder were abbreviated to wr. el. and sh. respectively. R and L refer to right and left side, respectively.

**Figure 6.**
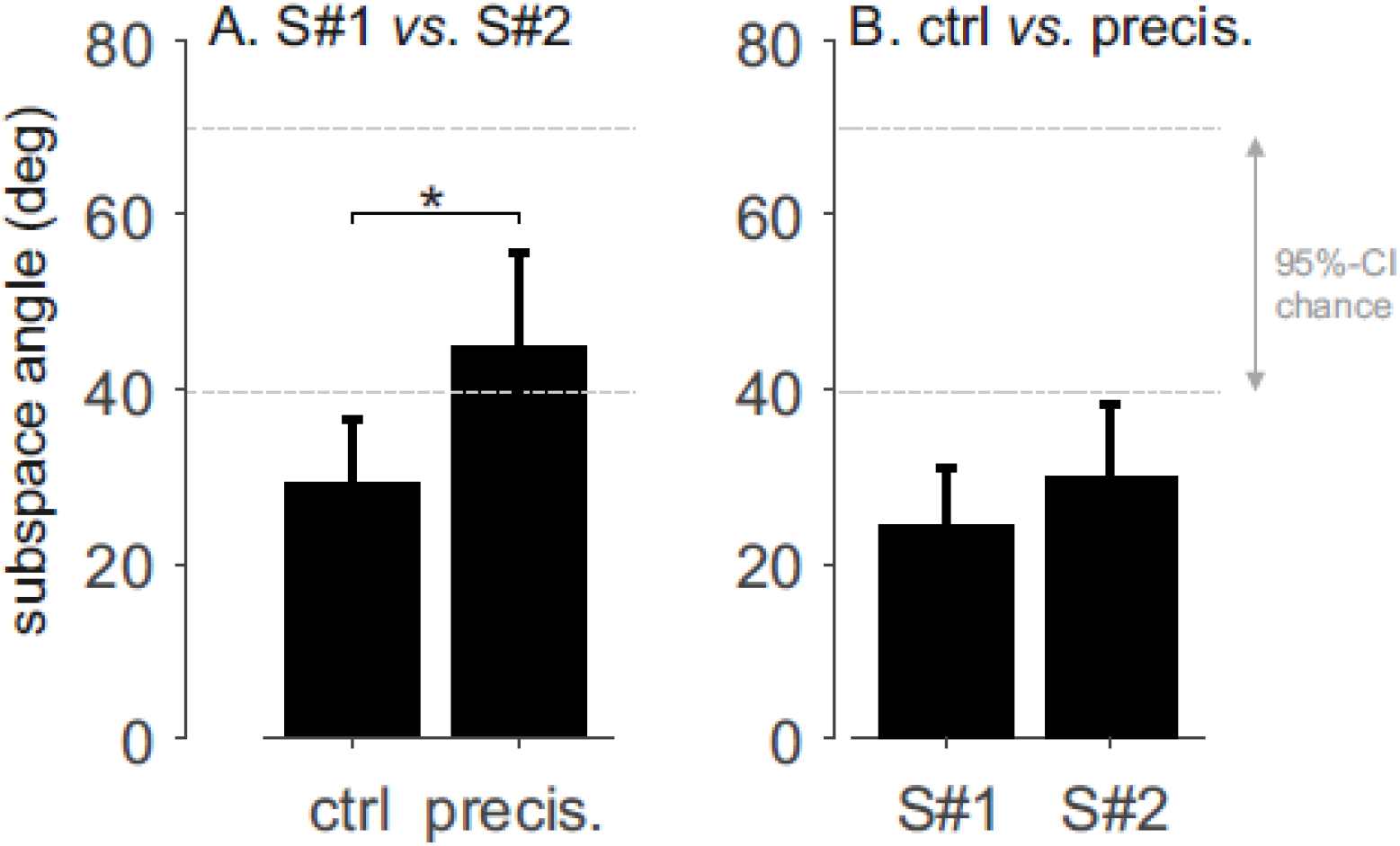
Subspaces comparison. The subspace angle measures the similarity between the subspaces spanned by the dynamic synergies. The 95%-confidence interval of angles obtained with random synergies (95%-CI chance) is indicated, i.e., CI=[49.5°,70.0°]. the star (*) indicates a significant difference (i.e., p<0.001).

For the conjoint PCA analysis, two dynamic synergies were extracted for all pairs of subjects, which accounted for 95.1±3.5% of total variance on average (range [84.4–98.8]%). The dynamic synergies for the conjoint analysis are depicted in Fig 7. VAFs were similar in the control and precision conditions (96.6±2.5% vs. 93.5±3.8%, respectively, Cohen’s d=0.7; Student t_9_=2.18; *p*-value=0.06). The subspace angles were not different than expected by chance (i.e., 46.4±21.3°compared to 36.3°, Student t_9_=1.50; *p*-value=0.17) when comparing the control and precision conditions (Fig. 7B).

**Figure 7.**
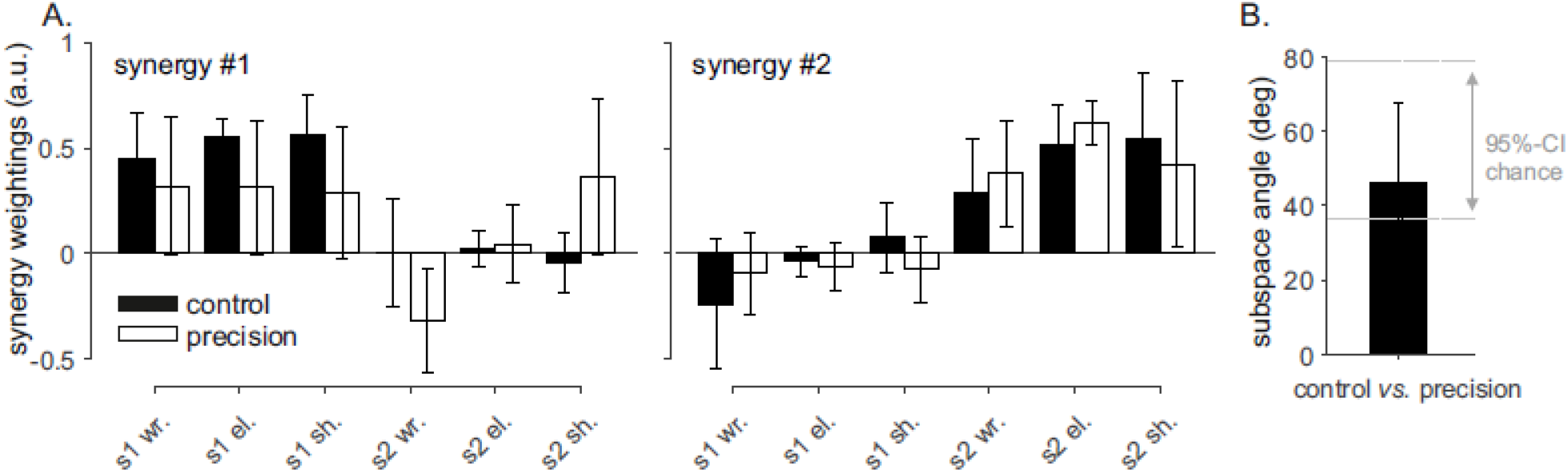
Conjoint synergies. S1 and S2 refer to participants #1 (right side) and participant #2 (left side), respectively. The 95%-confidence interval of angles obtained with random synergies (95%-CI chance) is indicated in panel B, i.e., CI=[36.3°,79.1°].

#### 4.1.4 Moment Cost Function (MCF)

In the CC the results obtained were divided into two different groups of dyads. Five dyads showed higher values of both Total *MCF* (Fig. 8A), (339,35 kg.m^2^.s^-2^ ± 20,44) and Δ*MCF* (Fig. 8B) while five other dyads showed lower values, (169,17 kg.m^2^.s^-2^ ± 17,06).

Regarding the PC, the results of Total and ΔMCF (respectively 172,66 kg.m^2^.s^-2^ ± 19,16; 5,78 kg.m^2^.s^-2^ ± 12,85) were less variable than in the CC.

### 4.2 Added task

#### 4.2.1 Accuracy score based on the ball trajectory

Regarding the ball trajectory, we found a correlation between the displacement of the handle and the ball on X-axis (Medio-lateral axis). The same goes for the displacement on the y-axis which represent the Postero-anterior axis (Fig. 4B). On the vertical Z-axis, the ball displacement and the handle are positively correlated.

Correlations were also computed in order to study the relationship between the ball trajectory and the sum of forces exerted by the subjects on the handles (Fig. 4C). Only two of ten dyads had a correlation on the X-axis; a positive correlation for the dyad 1 and a negative correlation for the dyad 10. On the Y-axis five of ten dyads had a significant negative correlation compared to the other three dyads who had a significant positive correlation. For the vertical Z-axis, nine of ten dyads had a positive significant correlation.

#### 4.2.2 Accuracy score

The mean Score_accuracy_ was 80.45±23.66 % during one gait cycle of the PACS in the PC condition.

### 4.3 Head and Trunk

Table 2 shows that when individuals had to keep the ball in the center of the target they turned the upper part of their body towards the box. Indeed, across the CC and the PC condition the orientation towards the box increased by 57.42, 9.22 and 3.29 degrees for the head, shoulders and pelvis, respectively. Also, the distance FOR-STE decreased by 7.69 cm between the CC and PC conditions, showing that the subjects were gazing at the box (Table S5).

**Table 2.** **Head, shoulders and pelvis orientation (angles in degrees) and distance between the forehead and the sternum (FOR-STE, in centimeters) in the CC and PC conditions;** mean (± s.d.). N=20 for each condition. * 0.05> p> 0.01; ** p <0.01 (paired Student t test).

|  | CC | S.D | PC | S.D |
| --- | --- | --- | --- | --- |
| Head orientation | 2.58 | ± 4.61 | 60.00 ** | ± 11.81 |
| Shoulders orientation | 1.99 | ± 3.13 | 11.21 ** | ± 6.61 |
| Pelvis orientation | 0.16 | ± 5.14 | 3.45 * | ± 4.05 |
| Distance FOR-STE | 21.67 | ± 4.86 | 13.98 ** | ± 3.25 |

## 5 Discussion

In this experiment, 10 dyads transported a load in two different conditions: a Control Condition (CC), in which they walked together while transporting a load, and a Precision Condition (PC), in which they walked together while transporting a load and maintaining at the same time a ball on its top. The first objective of our study was to test the hypothesis that the gait performance of two individuals walking together while transporting a load (CC) is disturbed when a precision task is added (PC).

We studied the center of mass of the system formed by the paired subjects and the box they carried. The result showed that the CoM_PACS_ speed decreased when the precision constraint was added. Besides, the second task induced a decrease in the pendular behavior and amplitude of the system. However, the period of the CoM_PACS_ displacement was not affected in PC. These results could be expected as the added task can be considered as a fine motor skill (Exner, 2001). Indeed, and as observed for the gait of an adult performing a dual-task (Yogev-Seligmann et al., 2010), the individuals needed to reduce their speed to perform both tasks. Similar to the findings of Holt et al., (2003) in a single carrier, the decrease in speed was accompanied by a decrease in the CoM_PACS_ vertical amplitude. These adaptations are reminiscent of the classic speed-accuracy tradeoff and the reduction in speed is very likely a strategy to reduce motor noise and improve controllability.

To explore the task further, we also studied each subject as a distinct entity. The comparison of the individual CoM trajectory of both subjects for CC and PC did not show any significant difference. Indeed, when walking side-by-side with a sensory interaction subject tend to synchronize their walking pace and kinematics (Schmidt and Turvey, 1995; Zivotofsky et al., 2012). The same goes for the CoM parameters (velocity, and CoM amplitude), i.e., no significant differences were found between the subjects at the left and the one at the right. The comparison of joint angles showed only a significant difference in hip angles between subjects.

No matter the condition they performed, the subjects tend to synchronize their speed and gesture frequency. However, the precision task altered the CoM_PACS_ kinematics leading to a less efficient energy transfer. In fact, this spatio-temporal strategy induced a decrease of the pendulum-like behavior at the CoM_PACS_.

Here, the energy recovered (*RR*) values obtained for the PACS in the CC (mean ± CI_0.95_= 60.25 ±8.57 %) were similar to those obtained in single carriers alone, as measured by Bastien et al. (2016) in Nepalese porters and in untrained individuals (*RR* = 61%), or by Tesio et al., (1998) in healthy individuals (*RR* = 60%). Our results showed a significant RR decrease in PC. This confirms the CoM_PACS_ pendulum-like behavior alteration. The potential and kinetic energy being out of phase, RR decrease leads to a higher mechanical cost of the whole system due to the precision task. On the other hand, the global muscular efforts estimated thanks to the moment cost function (MCF) of the upper-limbs were much more balanced between the individuals of each dyad when they performed the dual-task than in the lifting condition (Fig 8).

Doi et al., (2011) demonstrated that increasing the difficulty of a task (e.g., dual-task) affects the cost of the movement in elderly adults. Similarly, we found a modification in the trunk posture in PC that might have resulted in a finer control of the task. During PC, subjects spontaneously oriented their head, shoulders, and pelvis towards the box, probably to gather more visual information. Modification on the orientation of the upper part of the body seems to allow the subjects to look at the ball on the top of the load but, at the same time, these body segments were locked in a position that likely disturbed the kinematic of the PACS and his ability to behave as a pendulum. This interpretation is in accordance with the findings of Winter (1995), who suggested that during a bipedal walking, the control of the trunk restrains vision and head control. Here, the swings and rotations of these segments (head, shoulders, and pelvis), while locked during a gait cycle, do not contribute to the CoM_pacs_-evolution to time but may explain the lower-pendulum-like behavior in the PC. The decrease of the vertical amplitude of the CoM_PACS_ with both trunk and head fixed to look at the ball reveals a lower limbs pattern altered in PC. Numerous research studied the impact of the trunk posture on gait pattern, whether for medical purposes (Moraud et al., 2018) or sport performance purposes (Teng and Powers, 2014; Huang et al., 2019). These studies showed that a modification of trunk posture on the frontal and sagittal plane influences the bilateral lower limb kinematics and muscle activity. Here, the control of the walking speed and of the ball may have been supported by the lower legs and induced the dissipation of the mechanical energy thanks to eccentric work of the muscles.

The impact of the added task on the physical action of the subjects during the load transport was investigated. We recorded the forces applied by the subjects on the two box handles (Fig 4) during CC and PC. The subjects’ coordination was investigated through the correlation coefficient of applied forces. The results showed, in both conditions, a positive correlation for the forces on the antero-posterior and vertical axis. However, the correlation on the medio-lateral axis was weaker, with a large variability between the dyads of individuals. Thus, it seemed that the individuals coordinated their forces to move the load in the up-down and forward directions, without adopting a common strategy for the left-right direction. The results were similar between the two conditions, showing that the second task did not affect the collective strategies used during a simple load carriage task.

We considered that an action-reaction strategy was involved when the lag was higher than 150 ms, which corresponds to the minimum latency observed to take a decision after the perception of a stimulus (VanRullen and Thorpe, 2001). All lags were lower than 150 ms in the medio-lateral and antero-posterior axis. On the vertical axis however, lags higher than 150 ms were found in one dyad in the CC and in five dyads in the PC. Therefore, a modification of the behavior for half of the dyads was observed when the second task, requiring accuracy and precision, was added. These results might suggest a more conscious control of the box. It seems that the individuals moved the ball essentially by applying forces on the vertical axis at the handle inducing the rotation of the box around its anteroposterior axis and the displacement of the ball along the declination. When the box was thus moved by one individual, the second individual reacted (with a reaction time >150 ms) by moving it in the same direction in order to keep the ball in the center of the target.

Then, we used the moments applied on the box’s’ handles to compute through an inverse dynamic method the constraints at the joints of both upper-limbs of each subject. We observed a large variability of the joint moments among dyads in the CC. Half of the dyads produced much greater efforts than the other half and within this half, the joint moments produced by each individual were very unbalanced. In the precision condition, each individual within these five dyads produced similar efforts to keep the ball inside the circular target and to displace the load during a whole gait cycle. In both conditions, the participants were not allowed to communicate. However, in order to maintain the ball inside the target in the PC the dyad had to gaze at the box. Research on collective tasks showed that when sharing visual information’s of their performance, group members tend to coordinate their forces and movements (Bosga and Meulenbroek, 2007; Schmidt et al., 1998; Schmidt and Turvey, 1995). Hence, the visual feedback could have an impact on the muscular effort variability between dyads, as well as between individuals within each dyad.

These results suggest that the load carriage was affected by the second task, independently of the individuals’ performances in this task. Indeed, the accuracy score was not correlated (r=0.31, p=0.386) to the *RR*. The accuracy score was high, suggesting a good investment of the participants in the second task. According to Yogev-Seligmann et al., (2008) walking is a complex motor activity, which requires both the mobilization of executive functions, i.e. the cognitive capacities that allow an immediate adaptation of the motor behavior, and precision. On one hand, the decrease in locomotor performance in the precision condition could be explained by an increase in precision and a decrease in the mobilization of the executive functions used in locomotion. In the other hand, it also could be explained by a strategy of prioritization due to a structural interference between the precision needed to realize the first and the second task. Ebersbach et al., (1995), concluded that even when a task is highly practiced (e.g walking), adding concurrent tasks would lead to strategy changes depending on the attentional demand. Indeed, these control strategies are commonly used for humanoid robot when generating a movement prioritization. Sentis and Khatib, (2005), proposed a multi-level control hierarchy where the global task is decomposed into several subtasks. The hierarchy used ensured that constraints and critical task where accomplished first, while optimizing the execution of the global task. However, the absence of correlation between the accuracy score and the RR value reveals that the precision task may be too easy and may not discriminate different levels of precision.

Concerning the kinetic synergy analysis, a first observation is the non symmetry in terms of vector weightings for the loaded and unloaded arms i.e., the wrist, elbow and shoulder joint moments co-vary more with the neck and back joint moments for the loaded arm than for the unloaded arm, as demonstrated by high weightings coefficients for these joints (Fig 5). Subjects used more similar synergies during the CC than during the PC (Fig. 6A). The conjoint synergies were less similar than expected by chance when comparing the control and precision conditions (Fig. 7B). These two results show that a change in inter-joint moments coordination occurred due to the precision constraint. The synergies appeared more variable during the precision condition (Fig 5) and the weighting coefficients were “shared” between participants during the precision condition, e.g., the wrist joint moment for subject #2 was loaded with the wrist, elbow, and shoulder joint moment of subject #1 in the first conjoint synergy (Fig. 7A). These results suggest that although the coordination was more variable during the precision condition more co-variation occurred between the joint moments of the two participants. These results show that the collaboration during the precision task required disorganization of the spontaneous coordination adopted by the participants when no precision constraint was present. The change in posture between CC and PC might partly explain this observation. The VAF for the conjoint analysis tended to be lower during the precision condition (i.e., p=0.06) also suggesting a more variable coordination pattern between the joint moments of the two participants. These results suggest that coordinated action between two subjects does not necessarily require a similar coordination pattern for each of them, i.e., similar dynamic synergies, and that, in our experiments, a new coordination pattern emerged between the two subjects (i.e., different joint synergies), with more co-variation between their joint moments.

Our study highlights the fact that, when a dyad of individuals collectively transports a load and performs a second task (requiring accuracy and precision), the displacement of the CoM of the whole system (PACS) is affected, inducing a less efficient pendulum-like behavior. In both conditions the individuals coordinated their forces to move in the vertical and forward direction without adopting a common strategy in the left-right direction. We also observed that the individuals changed their trunk orientation and their behavior to manage the displacement of the ball inside the target. Furthermore, the visual feed-back permitted the dyad to coordinate their forces and movements in order to better control the position of the ball. However, the kinetic synergy analysis showed that subjects altered the structure of their own synergies in PC to adopt a coordination that was dissimilar between subjects but in which their wrist joint moments co-varied more. These results could be of interest for people working in ergonomics and could find potential developments in robotics (e.g. human-robot interactions), and in the rehabilitation domain, for example, when several caregivers in health care establishments have to move a patient.

## 6 List of symbols and abbreviations

CC: Control condition
CoM: Center of mass
CI: Confidence Interval
MCF: Moment Cost Function
PACS: Poly-Articulated Collective System
PC: Precision Condition
PCA: Pincipal Component Analysis
RR: Recovery Rate
VAF: Variance Accounted For

## 7 Author contributions

NS: responsible for the data analysis, interpretation and manuscript writing.GF: responsible for th edata analysis and interpretation, and major revisions of the manuscript. VF: responsible for the study design, supervision, data interpretation and major revisions of the manuscript. NAT: responsible for the data analysis and interpretation and major revisions of the manuscript. PM: Responsible for the study design, supervision, data interpretation and major revisions of the manuscript.

## 8 Competing interests

The authors do not have to disclose any financial or personal relationships with other people or organizations that could inappropriately influence (bias) their work.

## 9 Funding

This work was supported by the Financial support was provided by the Agence Nationale de la Recherche [CoBot-Projet-ANR-18-CE10-0003], the Association Nationale Recherche Technologie [CIFRE 2015/1321], and the MAS Marquiol for G.F. PhD grant.

## Notes

### Competing Interest Statement

The authors have declared no competing interest.

